## Supplementary Figures 1 -17 for "DeepLC introduces transfer learning for accurate LC retention time prediction and adaptation to substantially different modifications and setups"

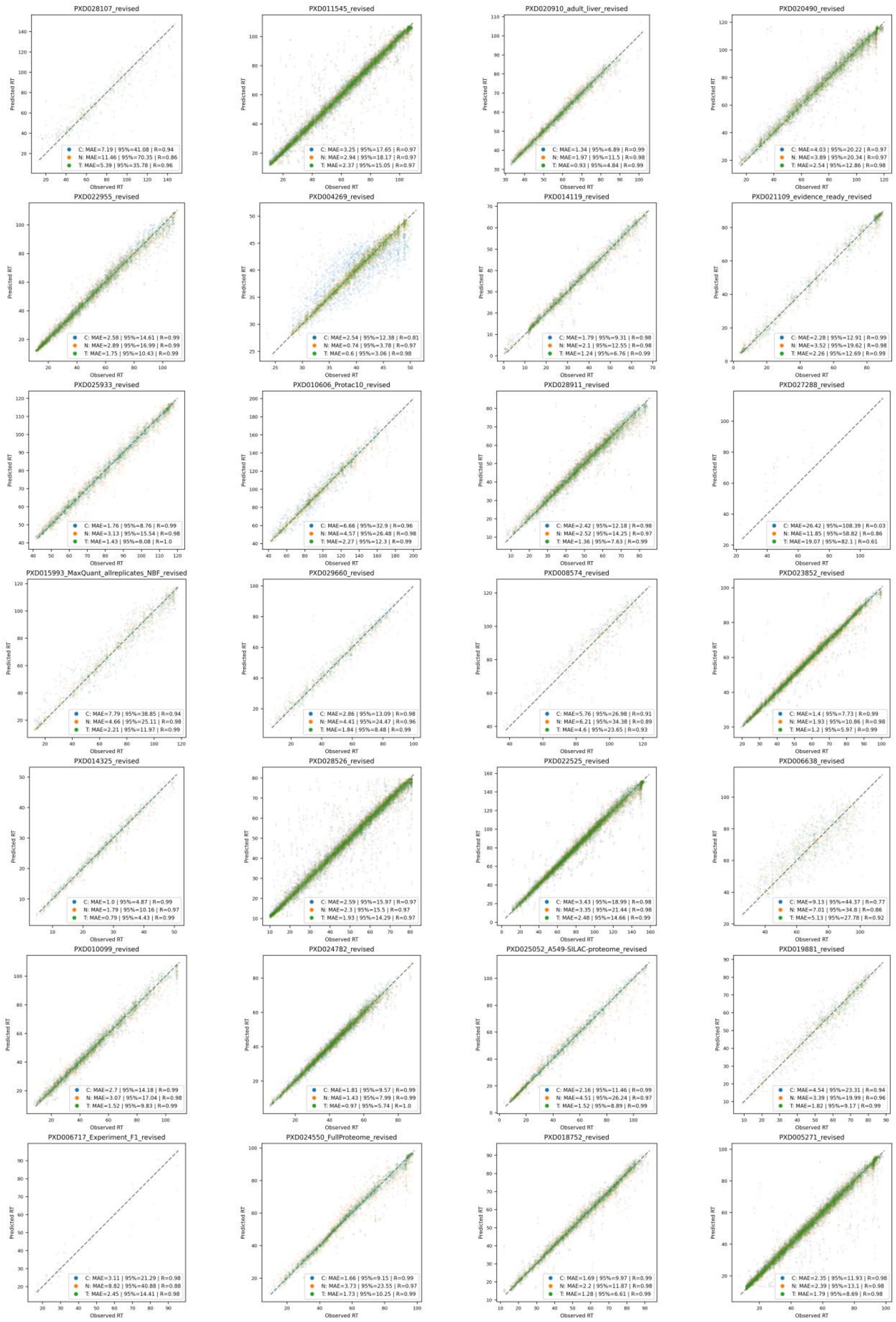


**Supplementary Figure 1.** Performance comparison of calibration (indicated with "C"), new model (random parameter initialization) (indicated with "N"), and transfer learning (indicated with "T") for 28 data sets.

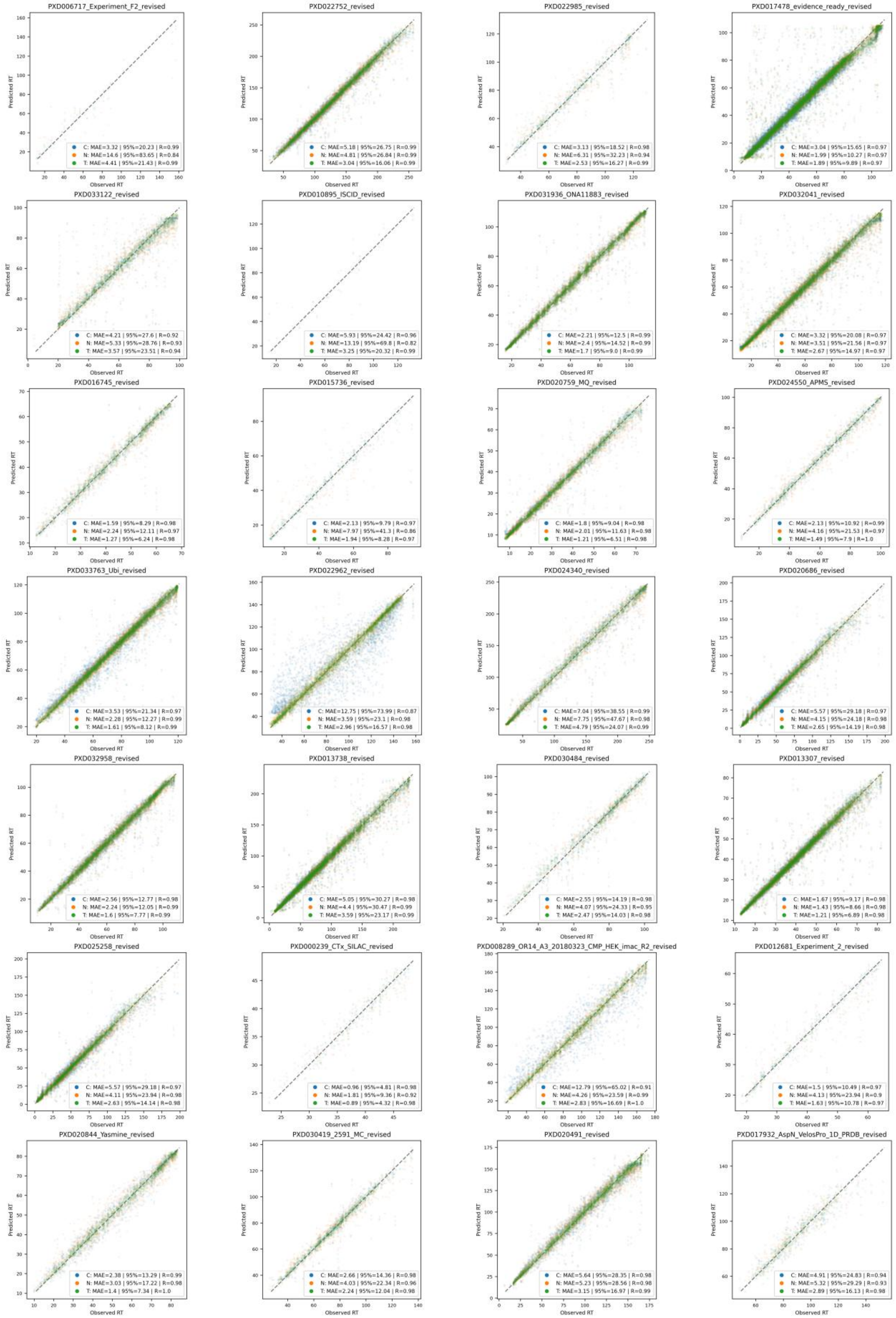


**Supplementary Figure 2.** Performance comparison of calibration (indicated with “C”), new model (random parameter initialization) (indicated with “N”), and transfer learning (indicated with “T”) for 28 data sets.

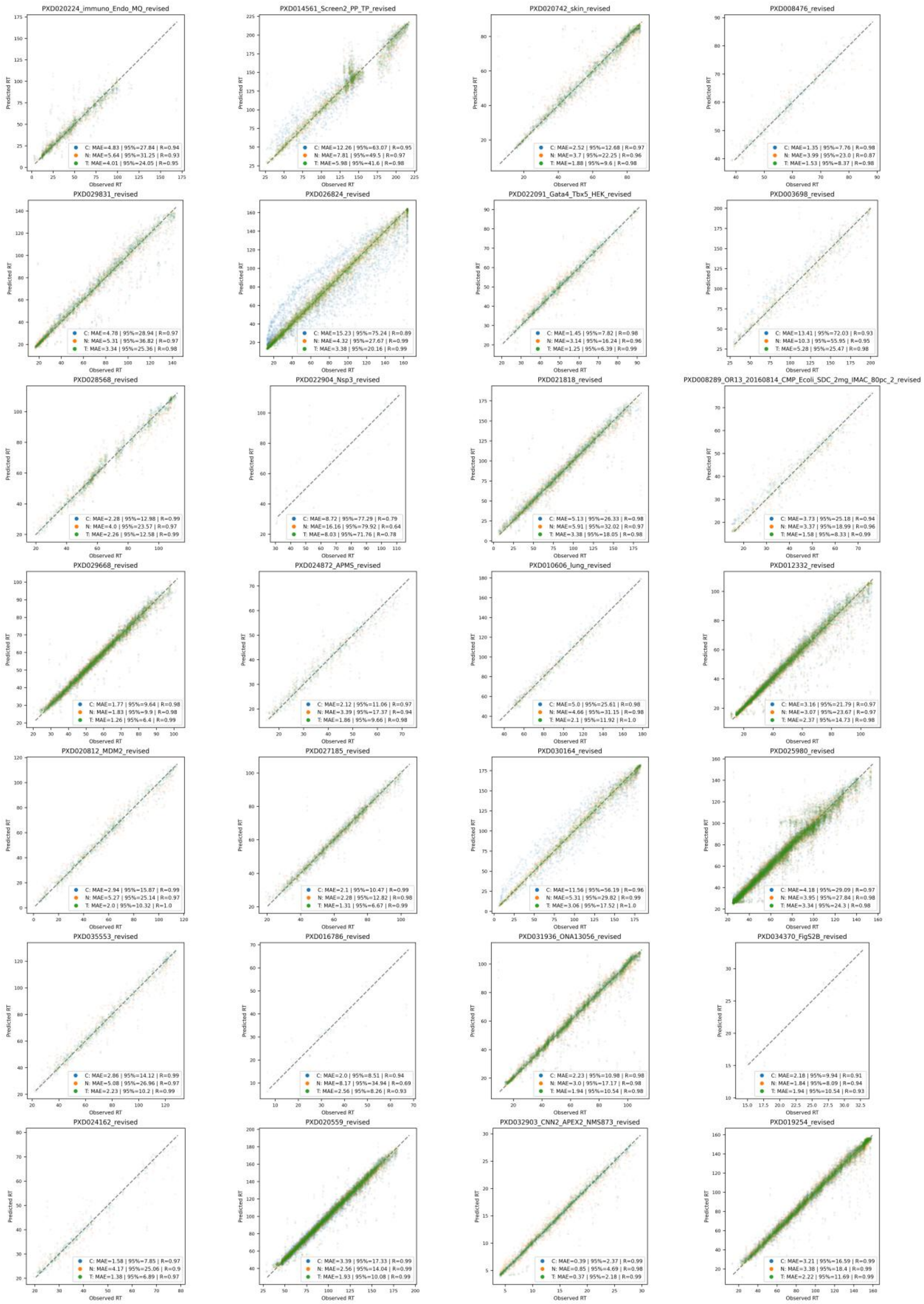


**Supplementary Figure 3.** Performance comparison of calibration (indicated with “C”), new model (random parameter initialization) (indicated with “N”), and transfer learning (indicated with “T”) for 28 data sets.

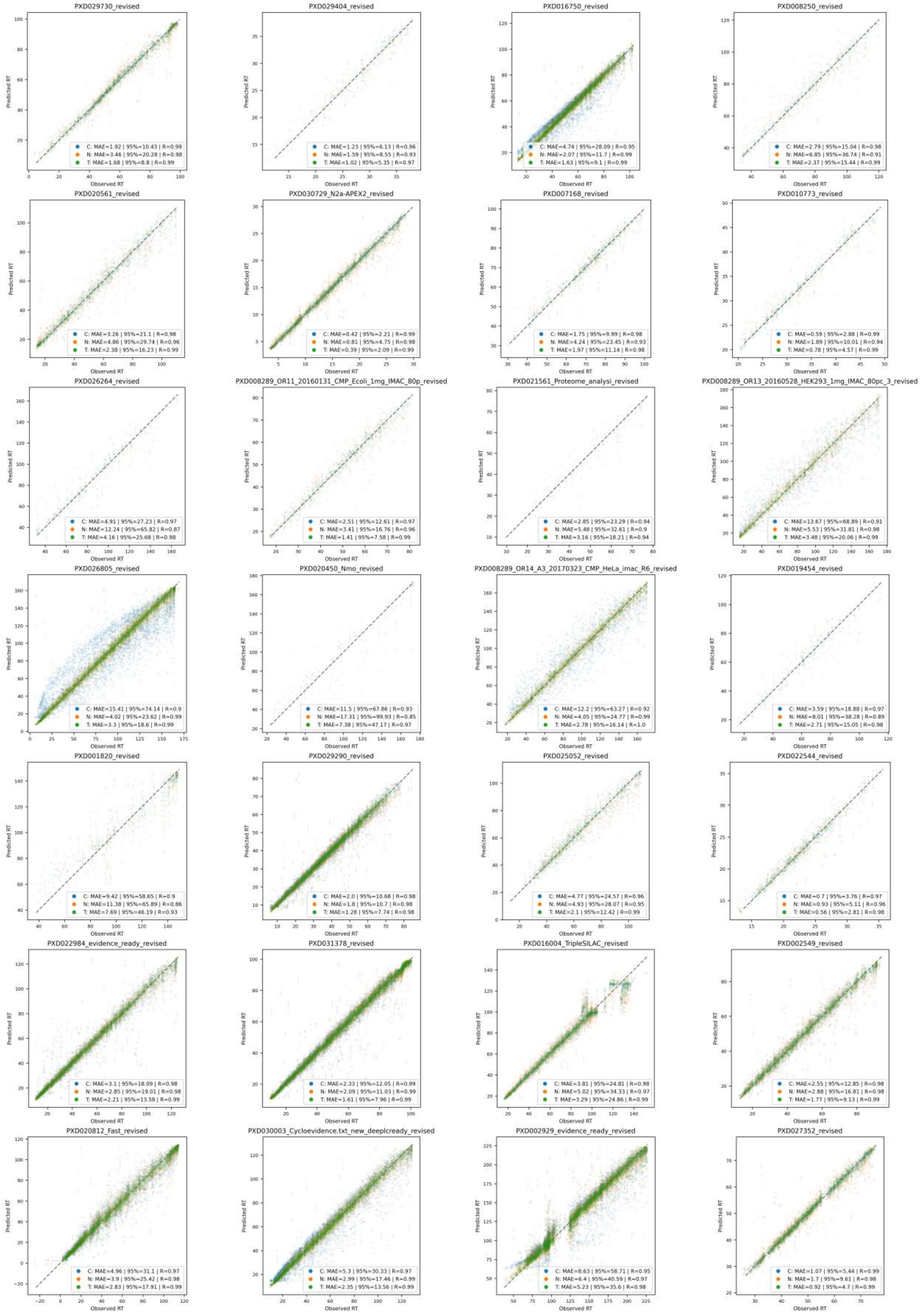


**Supplementary Figure 4.** Performance comparison of calibration (indicated with "C"), new model (random parameter initialization) (indicated with "N"), and transfer learning (indicated with "T") for 28 data sets.

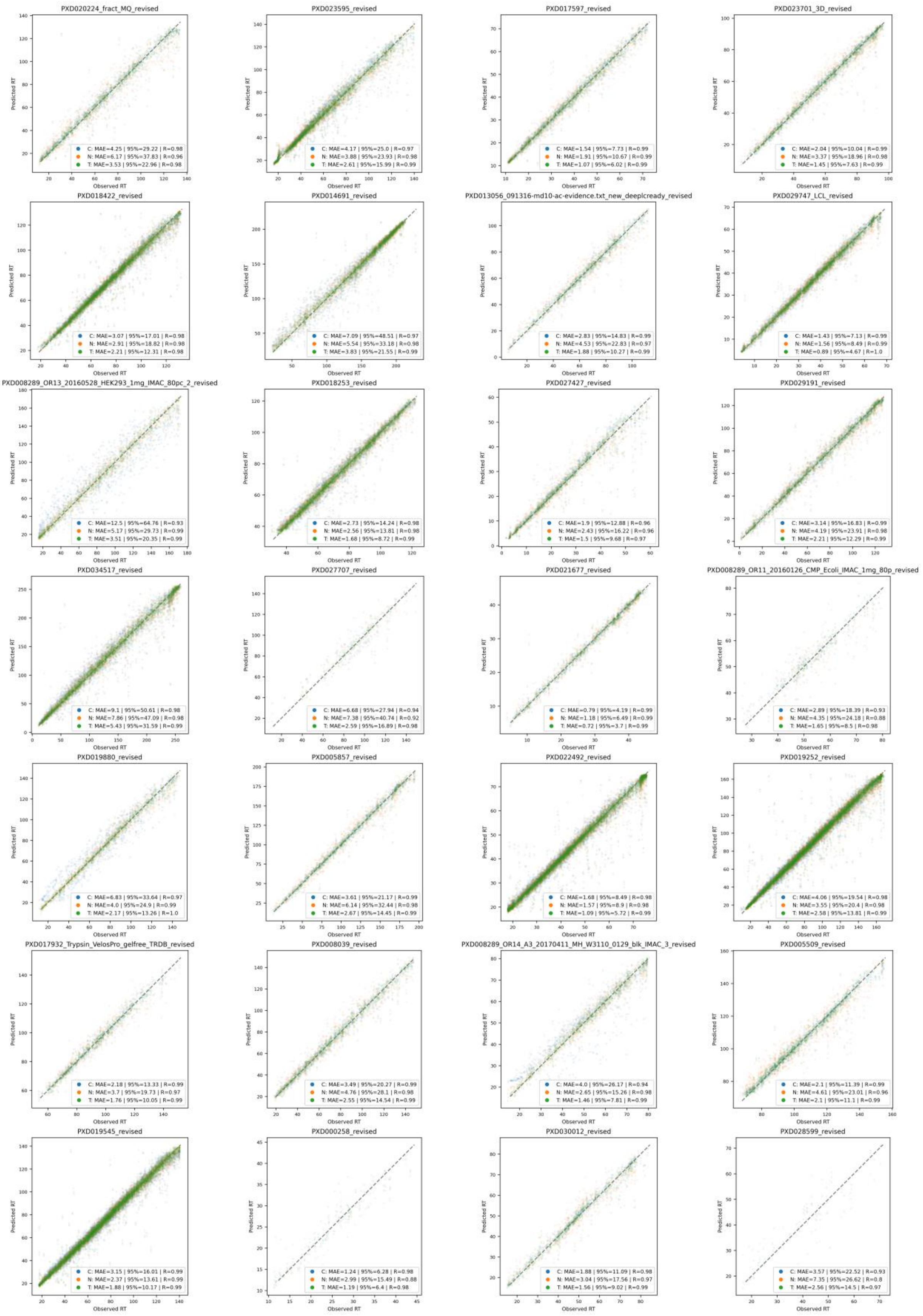


**Supplementary Figure 5.** Performance comparison of calibration (indicated with “C”), new model (random parameter initialization) (indicated with “N”), and transfer learning (indicated with “T”) for 28 data sets.

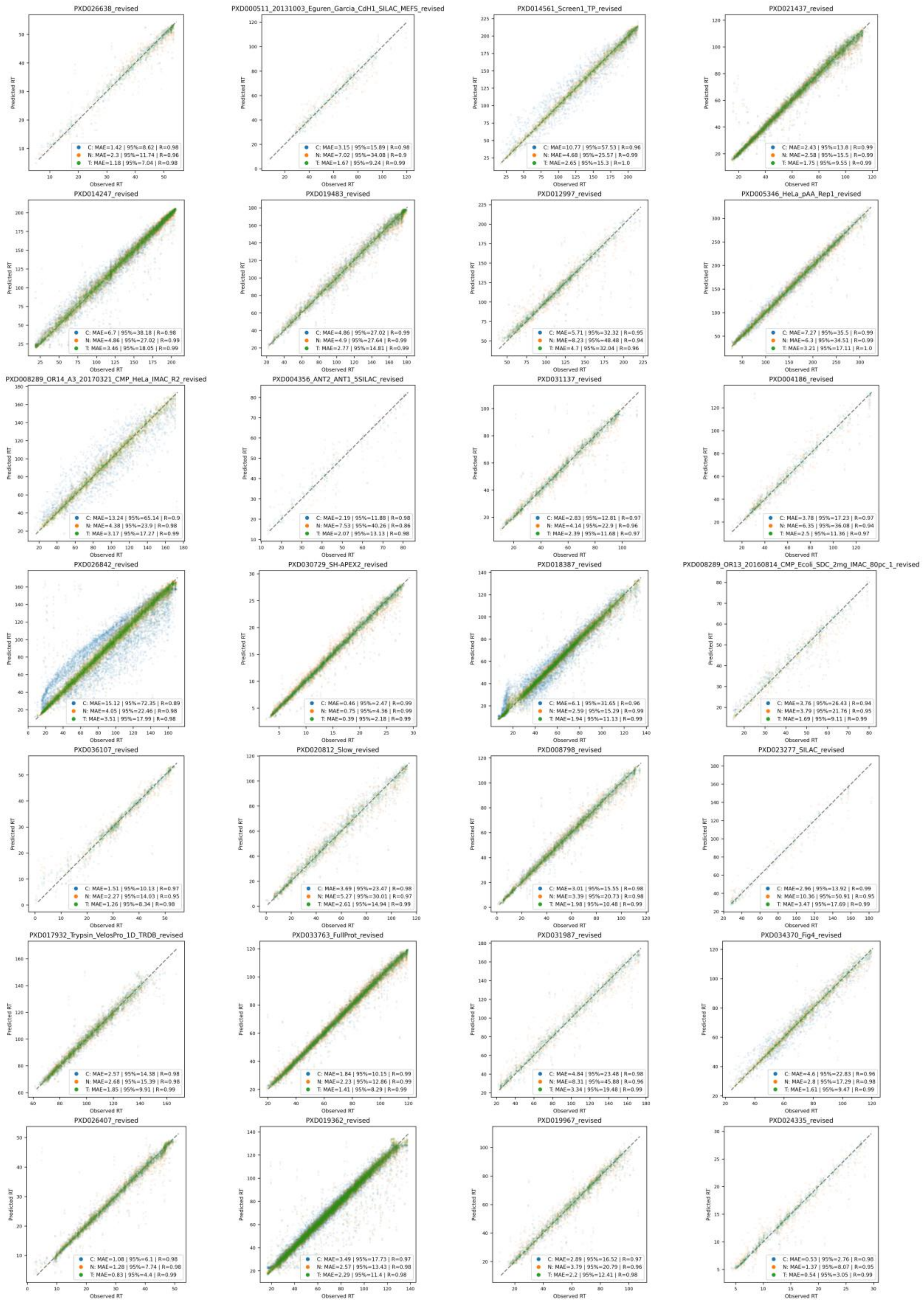


**Supplementary Figure 6.** Performance comparison of calibration (indicated with "C"), new model (random parameter initialization) (indicated with "N"), and transfer learning (indicated with "T") for 28 data sets.

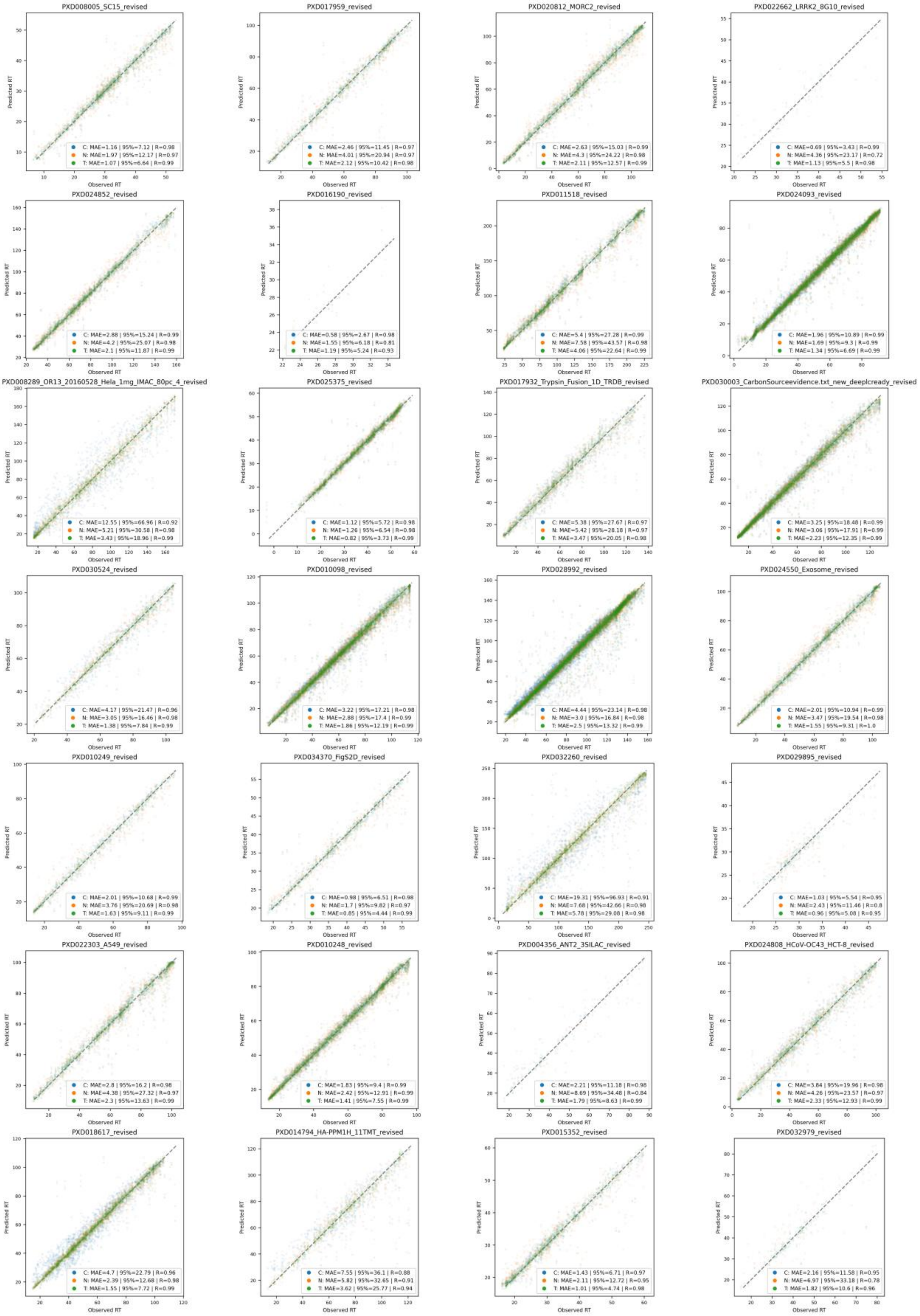


**Supplementary Figure 7.** Performance comparison of calibration (indicated with “C”), new model (random parameter initialization) (indicated with “N”), and transfer learning (indicated with “T”) for 28 data sets.

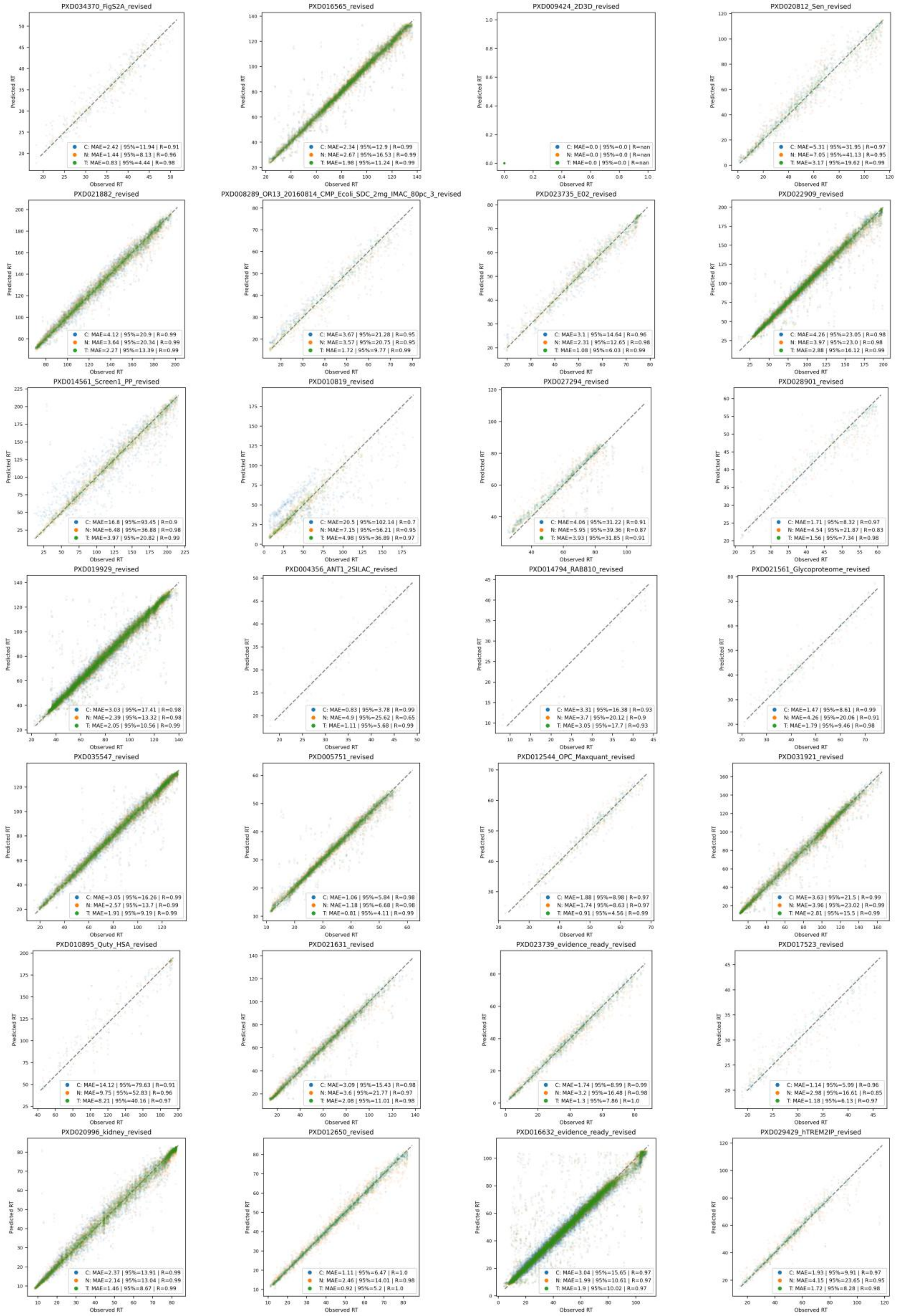


**Supplementary Figure 8.** Performance comparison of calibration (indicated with “C”), new model (random parameter initialization) (indicated with “N”), and transfer learning (indicated with “T”) for 28 data sets.

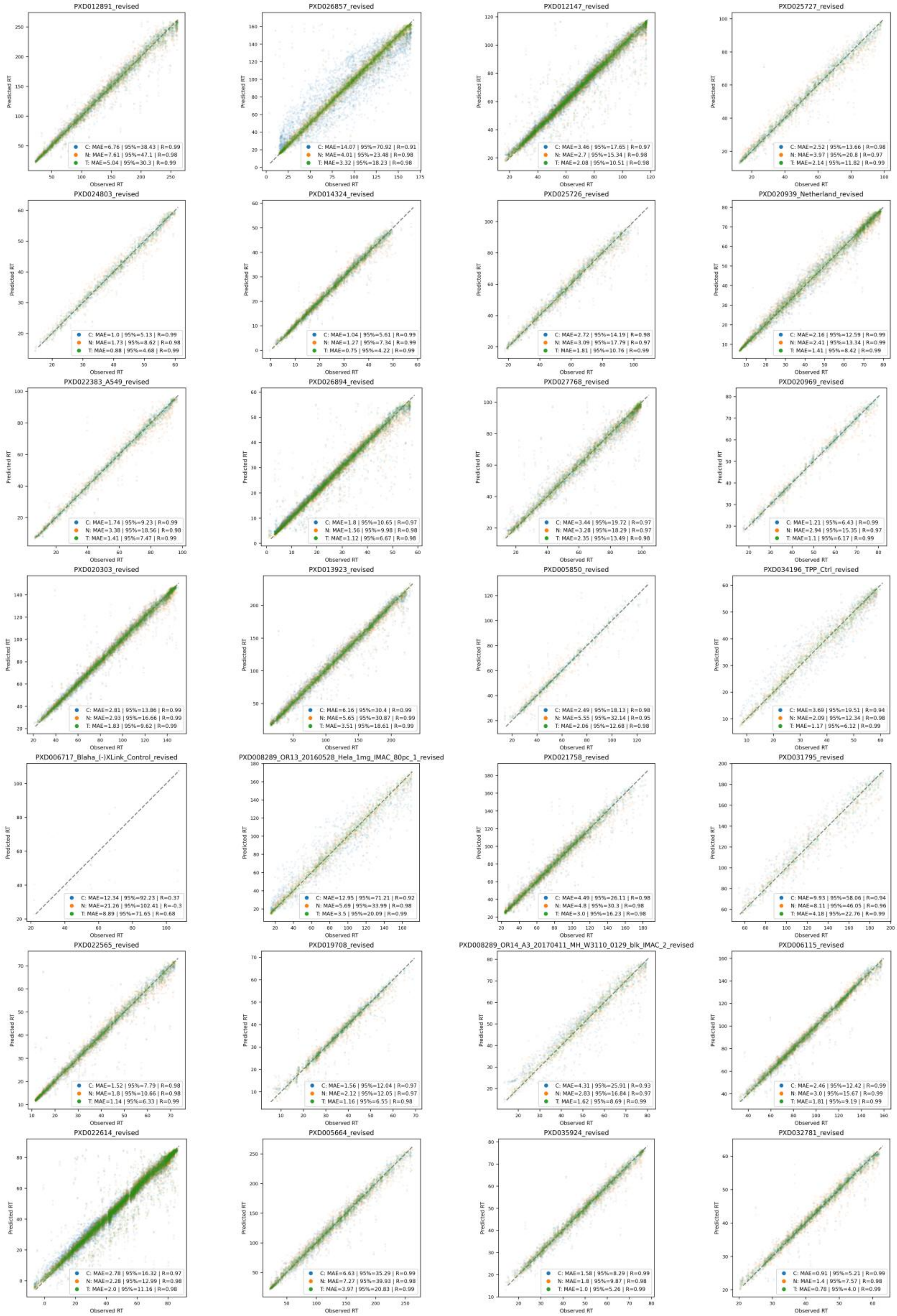


**Supplementary Figure 9.** Performance comparison of calibration (indicated with “C”), new model (random parameter initialization) (indicated with “N”), and transfer learning (indicated with “T”) for 28 data sets.

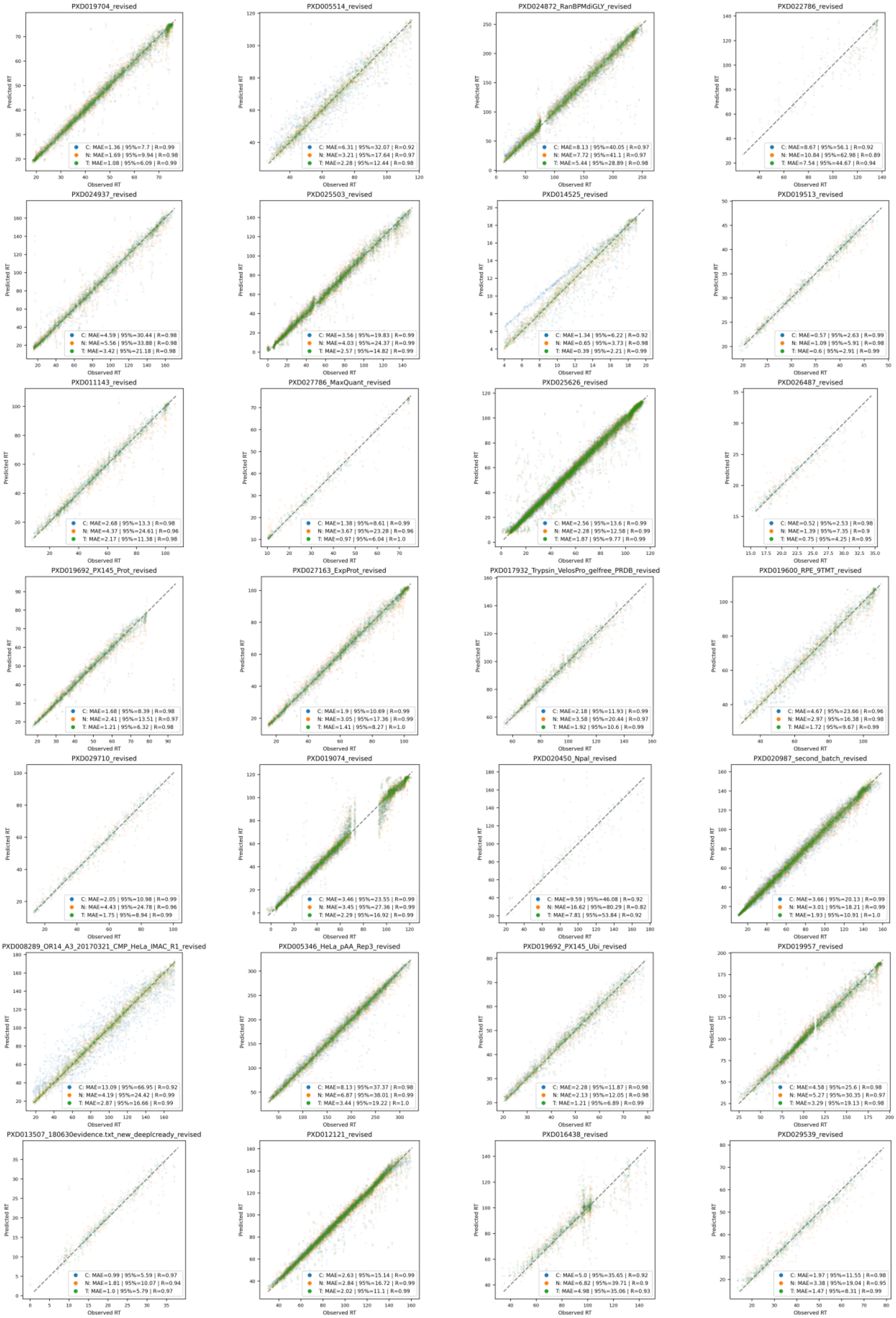


**Supplementary Figure 10.** Performance comparison of calibration (indicated with “C”), new model (random parameter initialization) (indicated with “N”), and transfer learning (indicated with “T”) for 28 data sets.

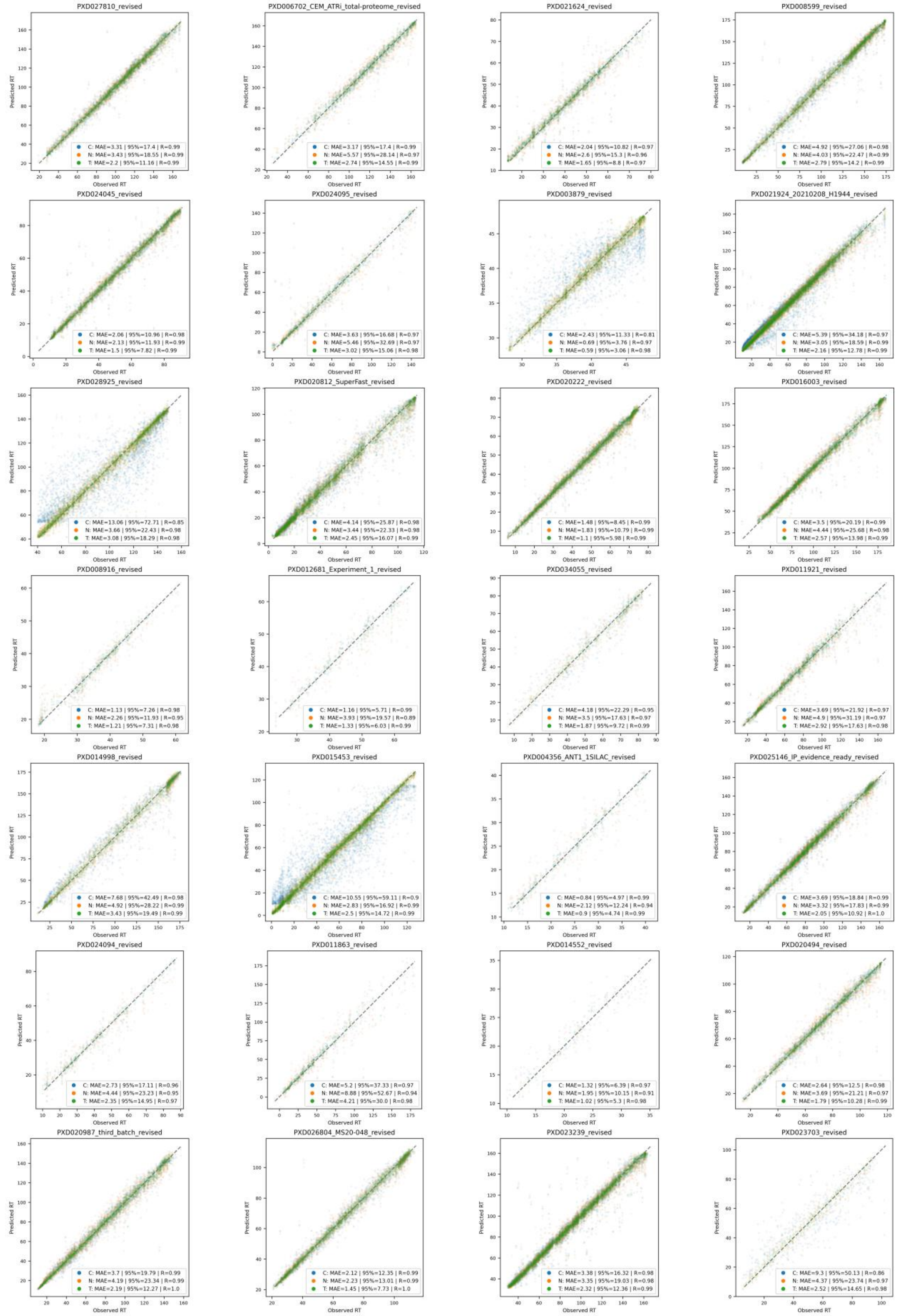


**Supplementary Figure 11.** Performance comparison of calibration (indicated with "C"), new model (random parameter initialization) (indicated with "N"), and transfer learning (indicated with "T") for 28 data sets.

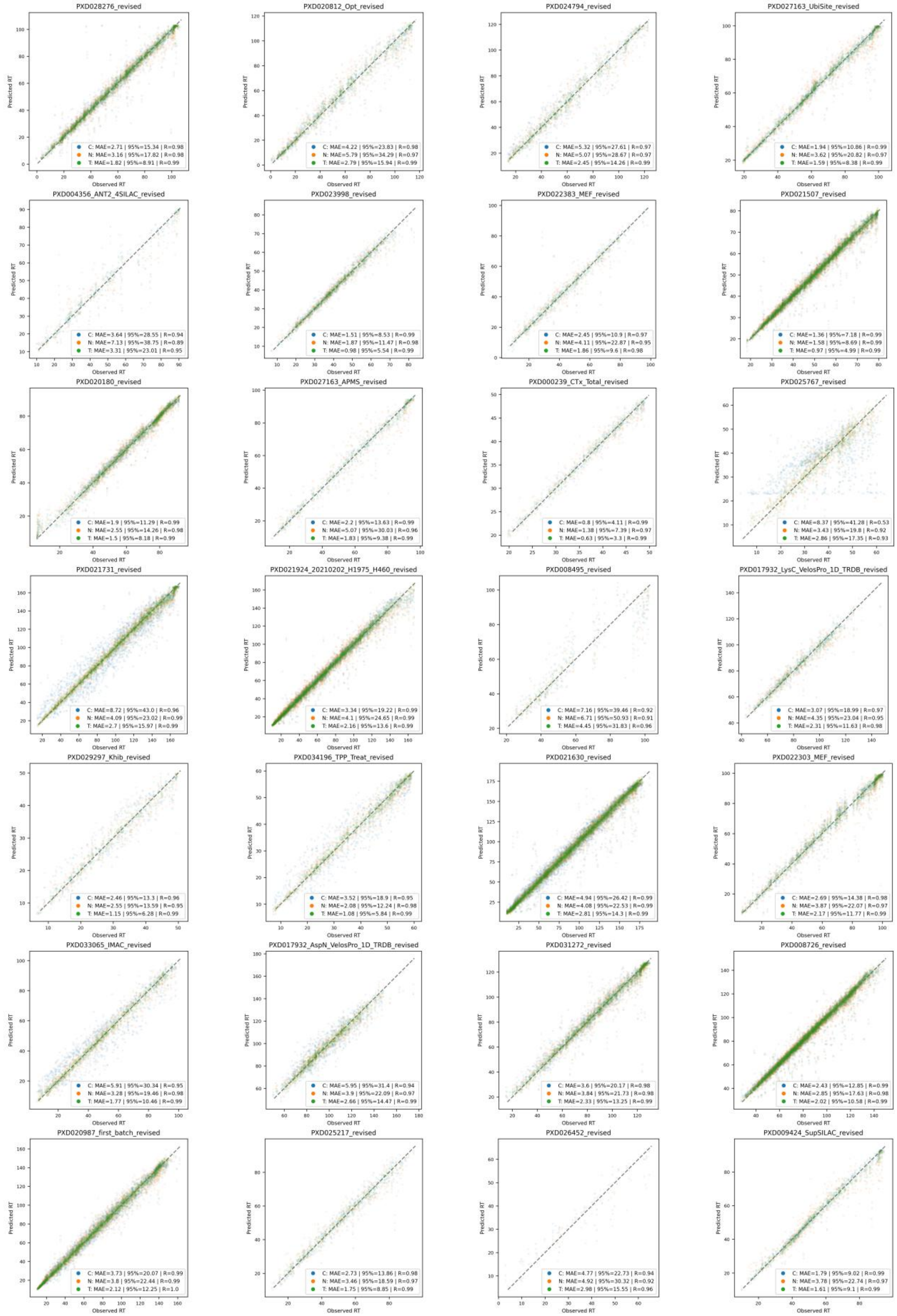


**Supplementary Figure 12.** Performance comparison of calibration (indicated with “C”), new model (random parameter initialization) (indicated with “N”), and transfer learning (indicated with “T”) for 28 data sets.

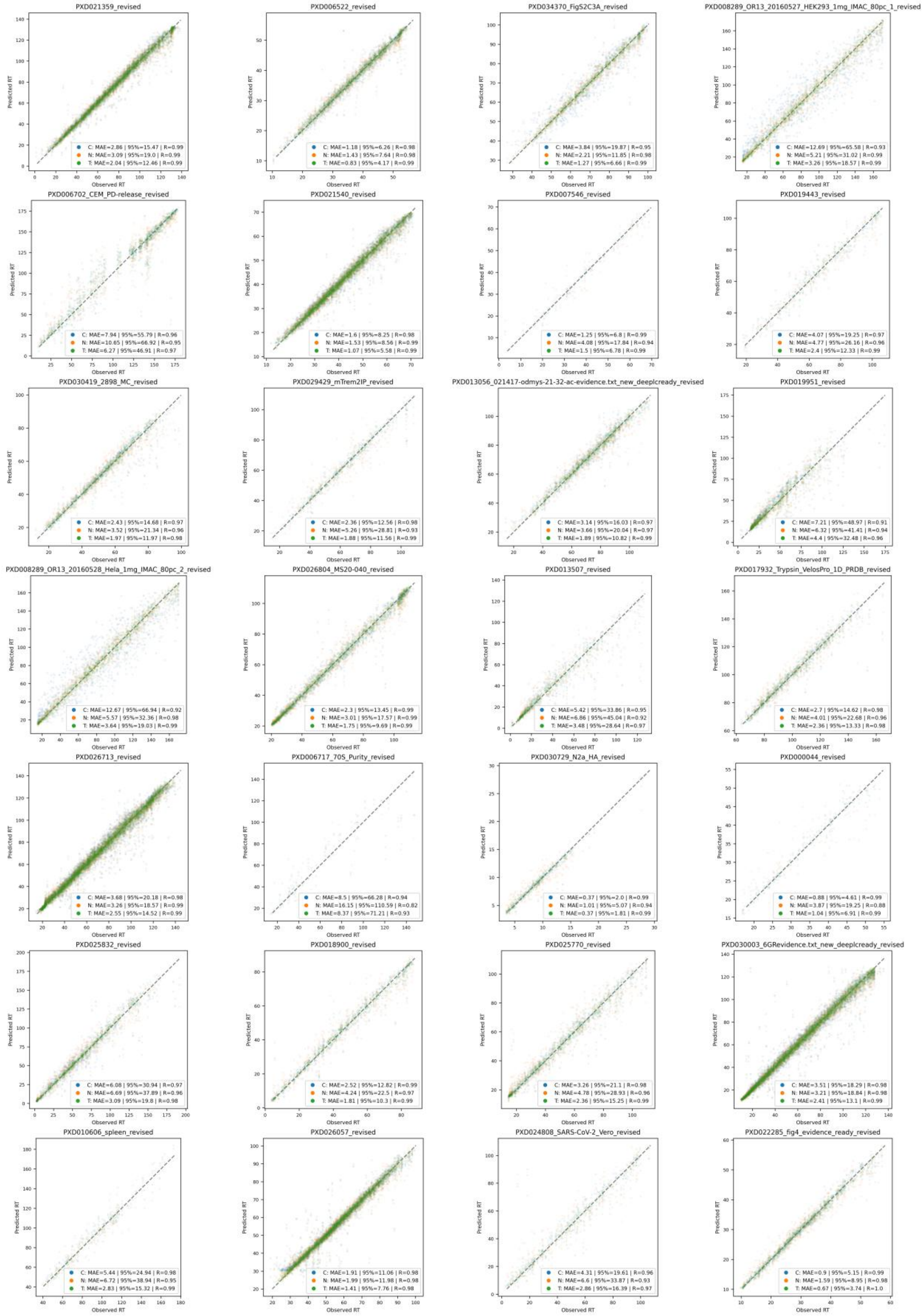


**Supplementary Figure 13.** Performance comparison of calibration (indicated with "C"), new model (random parameter initialization) (indicated with "N"), and transfer learning (indicated with "T") for 28 data sets.

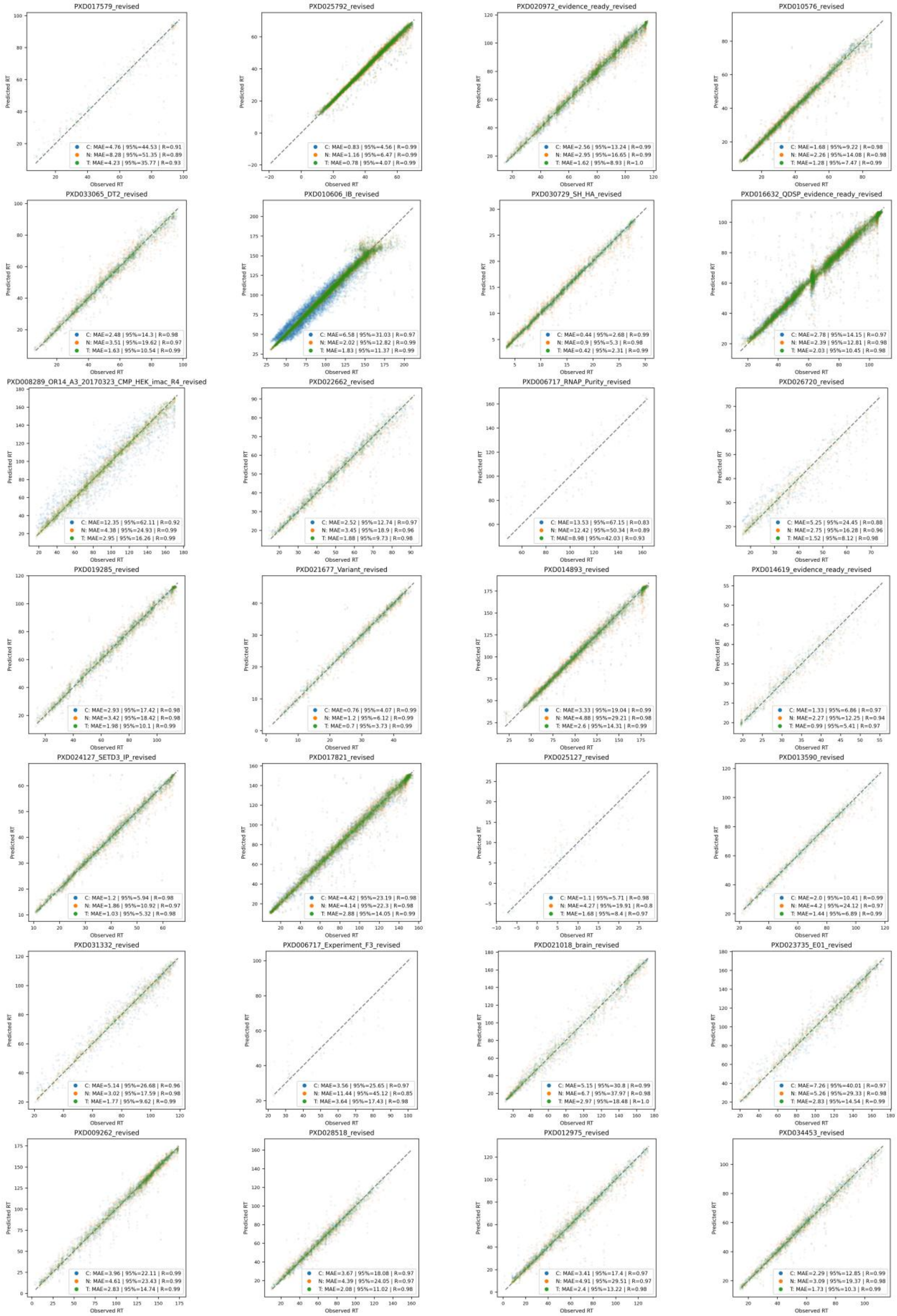


**Supplementary Figure 14.** Performance comparison of calibration (indicated with "C"), new model (random parameter initialization) (indicated with "N"), and transfer learning (indicated with "T") for 28 data sets.

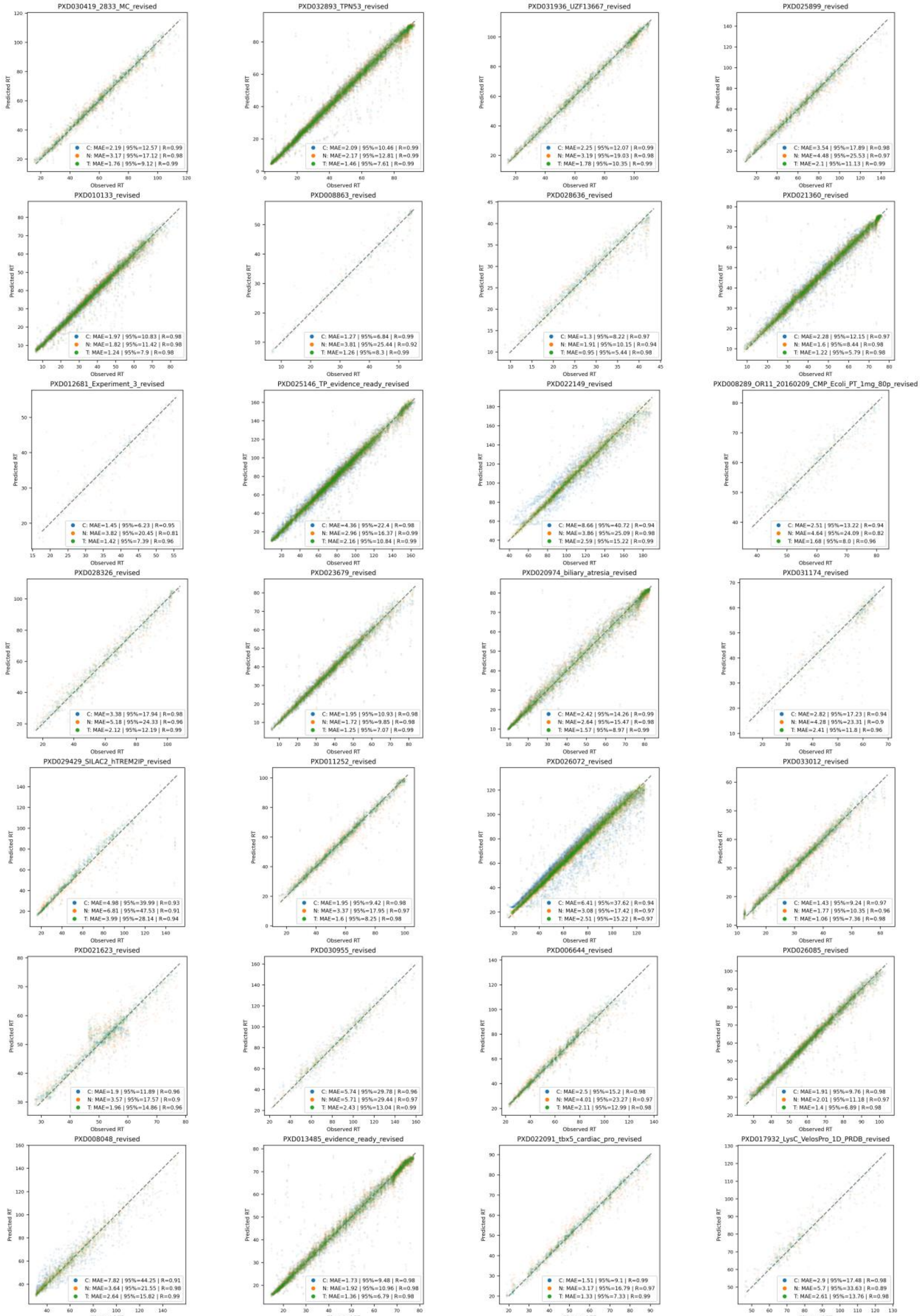


**Supplementary Figure 15.** Performance comparison of calibration (indicated with "C"), new model (random parameter initialization) (indicated with "N"), and transfer learning (indicated with "T") for 28 data sets.

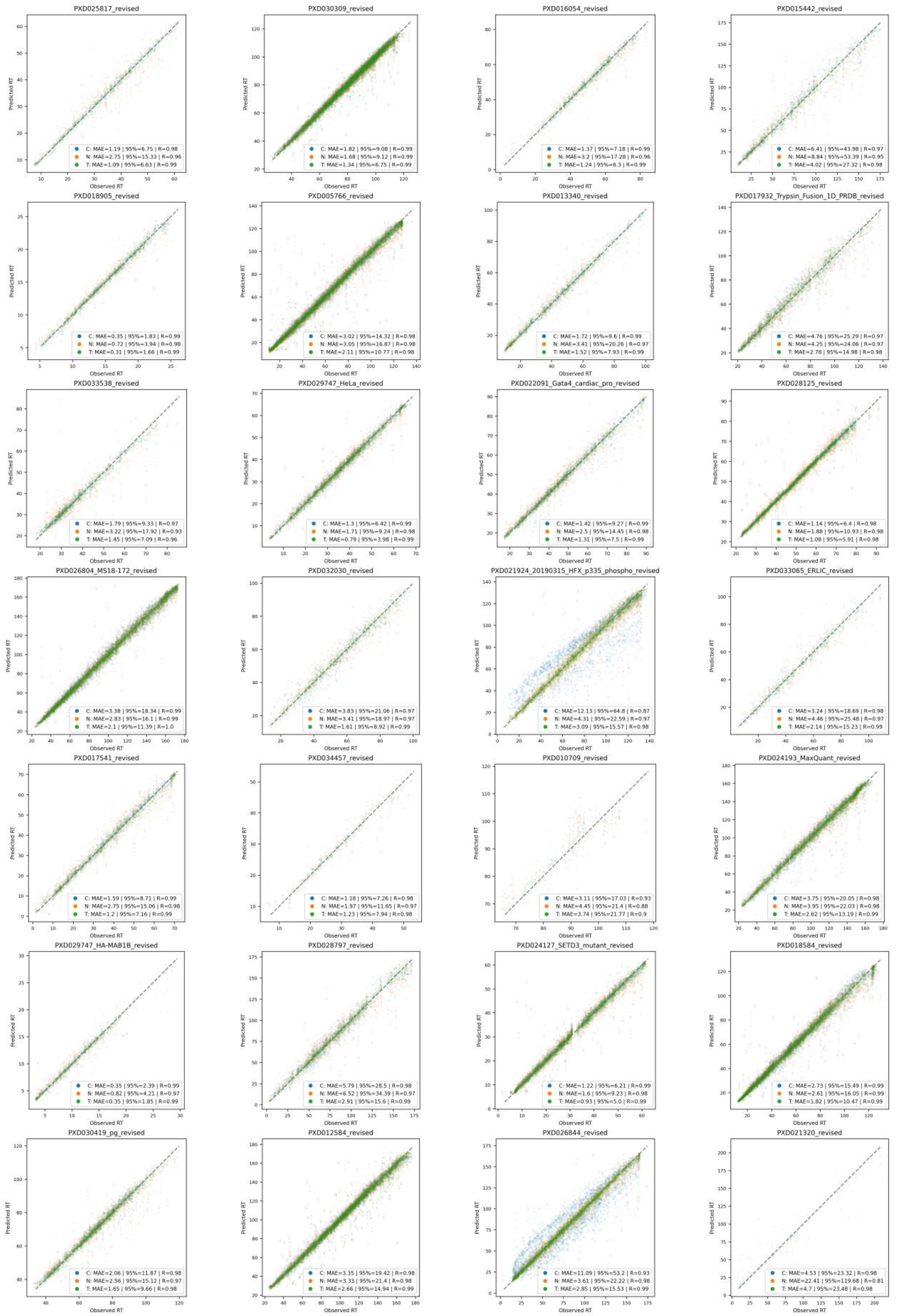


**Supplementary Figure 16.** Performance comparison of calibration (indicated with “C”), new model (random parameter initialization) (indicated with “N”), and transfer learning (indicated with “T”) for 28 data sets.

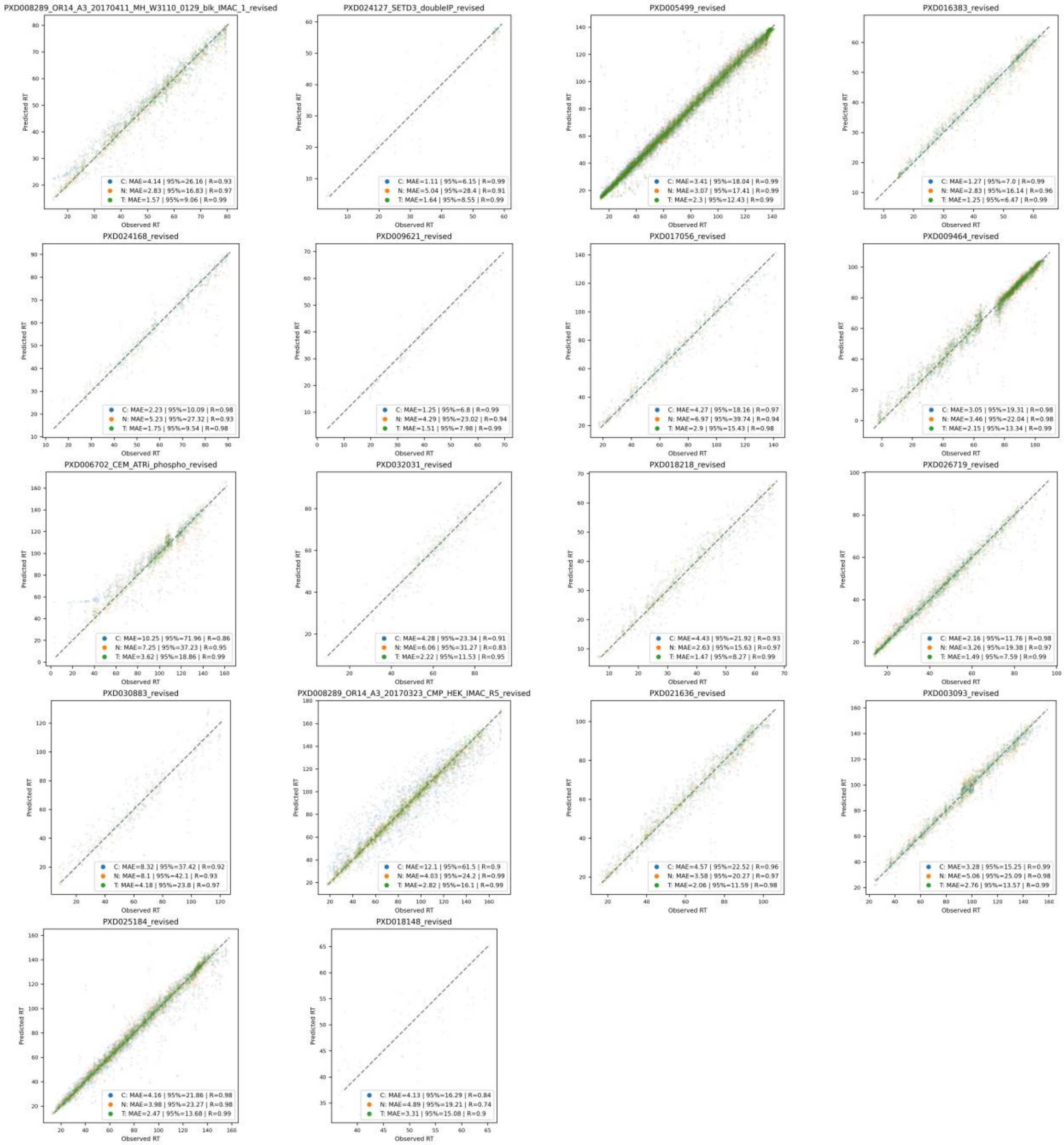


**Supplementary Figure 17.** Performance comparison of calibration (indicated with "C"), new model (random parameter initialization) (indicated with "N"), and transfer learning (indicated with "T") for 18 data sets.
